# Detection of low numbers of bacterial cells in pharmaceutical drug product using Raman Spectroscopy and PLS-DA multivariate analysis

**DOI:** 10.1101/2022.04.26.489535

**Authors:** R.A. Grosso, A.R. Walther, E. Brunbech, A. Sørensen, B. Schebye, K.E. Olsen, H. Qu, M.A.B. Hedegaard, E. C. Arnspang

## Abstract

Sterility testing is a laborious and slow process to detect contaminants present in drug products. Raman spectroscopy is a promising label-free tool to detect microorganisms and thus gaining relevance as future alternative culture-free method for sterility testing in pharmaceutical industry. However, reaching detection limits similar to standard procedures while keeping a high accuracy remains challenging, due to weak bacterial Raman signal. In this work, we show a new non-invasive approach focusing on detect different bacteria in concentrations below 100 CFU/ml within drug product containers using Raman spectroscopy and multivariate data analysis. Even though Raman spectra form drug product with and without bacteria are similar, a partial least squared discriminant analysis (PLS-DA) model shows great performance to distinguish samples with bacteria contaminants in limits below 10 CFU/ml. We use spiked samples with bacteria spores for independent validation achieving a detection accuracy of 99%. Our results indicate a great potential of this rapid, and cost-effective approach to be use in quality control of pharmaceutical industry.

## Introduction

Testing for microbial contamination is a crucial step in quality control of pharmaceutical drug products (DP) before their commercial release ^1, 2^. Standard procedures for bioburden testing are highly time-consuming, costly and are limited in terms of sensitivity and specificity given that they depend on growing conditions. ^3, 4^. Hence, the pharma industry needs to develop and implement faster and cost-effective novel technology for this purpose. Bacterial contamination criteria vary depending on the type of pharmaceutical formulations and administration route ^5, 6^. The complete absence of micro-organisms is required for quality control tests of drug products required to be sterile ^7^. For this reason, novel biotechnological approaches for contamination tests must secure detection of all microorganisms potentially present in the final product ^2, 8, 9^. Analyzing for absence of all micro-organisms by traditional growth based microbiological methods presents a further challenge for the laboratory environment to minimize the risk of false positive test results caused by laboratory contaminations during sample handling ^6^. Within the last decades new set ups involving spectroscopic technology have been studied for faster and simpler microbial detection and quantification in drug products ^9-12^.

Raman spectroscopy (RS) is a non-invasive method based on inelastic scattering of monochromatic light upon interaction with chemical bonds present in the sample. Each molecule gives a specific Raman spectrum (fingerprint) depending on their chemical environment, and chemical and biophysical properties ^13^. In combination with chemometrics, RS has been gaining significant consideration in the pharmaceutical industry given its reduced cost, faster quantitative analysis, and real-time monitoring of various processes that involve changes of the molecules and/or their chemical environments ^14, 15^. Concerning microbial contamination, RS has shown great performance detecting a wide range of bacterial components i.e., lipids, proteins, amino acids, nucleic acids ^16^. This technique is even able to distinguish among several bacterial strains present in water-based formulations and in solid drugs with high accuracy ^17, 18^. However, studies with RS showing high robustness while retaining the ability to distinguish even bacterial strains lack sensitivity and do not reach a sufficiently low limit of detection (LOD) of bacteria cell number for bioburden test acceptance ^19, 20^. Contrary, more sophisticated RS set ups with LOD below 10^3^ CFU/ml have demonstrated drawbacks in terms of reproducibility, data analysis and expensive technology making these methods inconsistent and difficult to scale up for pharmaceutical industry application ^21, 22^.

Even though advances with RS are promising, new approaches must improve its accuracy and reproducibility to be suitable for pharmaceutical industry quality control, and act as simpler, faster, and more cost-effective applications. Remanent challenges when detecting bacteria with RS in pharma products are (i) discrimination between Raman spectra from organic molecules present in the formula and bacterial ones, (ii) detection at low contamination given the weak signal from the bacteria in comparison with the product volume (iii) contribution to the Raman signal from other sources such as product packaging, fluorescent compounds, and (iv) correct data processing and statistical analysis model ^14^. In this study, we present a novel approach to up-concentrate and detect ≤10 CFU/ml of relevant bacteria with RS and multivariate analysis without breaching the primary DP package. The outcomes of this project support RS as non-invasive and non-destructive method to detect bacterial contamination in DP as alternative to the pharmacopeial ^7^ destructive method susceptible to laboratory contaminations. Furthermore, the developed method has the potential to enable real-time monitoring of the contaminations in pharmaceutical processing.

### Experimental

#### Reagents and bacteria strain

The pharmaceutical product (DP) contains 3 ml of solution with 100 U/ml insulin, glycerol, 1.80 mg/ml phenol (CAS 108-95-2), 2.06 mg/ml m-cresol (CAS 108-39-4) (preservative), zinc acetate, glycerol, phosphate buffer, sodium chloride and water for injections. The primary package is a vial made of 2 mm borosilicate glass.

Freeze-dried preparations in BioBall^®^ MultiShot 550 of *B. subtilis* (NCTC 10400) and *S. enterica* (ACM 5080) were from bioMérieux. *B. subtilis* in spore form. *S. haemolyticus* (ATCC 29970) was kindly provided from Microbial Competence Centre, Novo Nordisk. Other reagents were Ethanol (20C184005, VRW chemicals), Phosphate buffered saline solution (S3308, MP Biomedicals), Luria-Bertani broth agar (SLCC1516, Sigma) and Tryptic soy broth (TSB) (ICNA091010717, MP Biomedicals). Tryptic soy agar (TSA) was prepared by adding 20 g/L of agar (Sigma) to TSB solution.

#### Bacterial culture and growth conditions

For experiments with vegetative forms of *B. subtilis*, one BioBall^®^ MultiShot 550 of *B. subtilis* was inoculated in LB broth media and incubated at 35 °C for 18-24 hour and then sub-cultured in LB agar and incubated in aerobic conditions at 30-35 °C for 24 h. Bacterial numbers were estimated by colony harvesting, dilutions in sterile PBS and counting in chamber under the microscope, then applying dilution factor calculation. Experiments with low CFU number (50 and 10 CFU) were performed directly with dilutions from the dissolved BioBalls (550 CFU/pellet). *S. enterica* and *S. haemolyticus* were cultured using TSA plates. Experiments with low numbers of *B. subtilis* spores were made by dilutions from BioBall^®^ MultiShot 550 of this bacterium and then spiking the samples with the respective amount to reach a final concentration of 50 and 10 CFU/ml. Controls of injected CFU number were made by enumeration in TSA plates and direct counting of colonies.

#### Sample preparation and Raman spectroscopy

Spiked vials for experiments with 3 × 10^8^, 50 and ≤10 CFU/ml of bacteria were prepared using the following procedure. Briefly, in a laminar air flow (Heraguard ECO 0.9, Thermo Scientific), bacterial dilution was prepared using sterile DP solution as diluent and homogenized by vortexing. Then the precise volume was injected into the products vials (samples) using a calibrated syringe (Hamilton^®^ syringe 1700 series) to get the desired CFU concentration. The vials were placed in a self-built plate and centrifuged at 2500 rpm for 17 minutes. Before experiments, vials’ outer glass surface was cleaned with ethanol avoiding any Raman signal from organic material adsorbed during sample handling. Given that the DP contains preservatives, a parallel control was run covering the whole procedure time from CFU injection till last sample measurement with RS. This control was cultured along with analyzed sample to evaluate bacteria survival without exposure to laser.

Raman spectra were recorded at room temperature using a multimode diode laser (FATBOY model, Ocean Insight) emitting at a wavelength of 785 nm. The laser excitation was fiber-coupled to a focusing 785 nm immersion Raman probe (Ocean Insight) with a 5 mm focal length delivering a total laser power of 350 mW at the sample. The Raman back-scattered light collected by the probe was directed through a low OH multimode fiber to the spectrometer (QE Pro Raman Series, Ocean Insight) equipped with a charge coupled device operated at -50 °C. For each sample, 10 Raman spectra were acquired with an integration time of 10 seconds. Spectrometer setup, data acquisition and control were done with Ocean View 2.0.8 software (Ocean Insight).

### Data processing and analysis

Raman spectra were pre-processed using MATLAB R2021a software (MathWorks, US). Data was truncated to the wavenumber region 400-3200 cm^-1^, baseline corrected using asymmetric least squared smoothing ^23^, normalized (area under curve) and mean centered. PLS-DA ^24^ was performed using the PLS-regression algorithm in MATLAB. For model calibration, we used a total of 100 spectra of sterile and 100 spectra of vegetative *B. Subtilis*. For experiments with *S. enterica, S. haemolyticus*, and *B. subtilis* spores a total of 80 Raman spectra from sterile and spiked samples were analyzed. To differentiate *B. Subtilis, S. haemolyticus* and *S. enterica* from sterile conditions a 10-fold cross validated PLS-DA model was applied to each case with 50 and 10 CFU/ml. For *B. Subtilis* spores, *S. haemolyticus* and *S. enterica* four, three and three components were used in the model respectively based on the results of the cross validation **(see Table 1)**. An independent validation set of spectra from *B. Subtilis* spores was tested against the corresponding model calibration to obtain the specificity and sensitivity of the model.

**Table 1:** Values of root-mean squared error of calibration (RMSEC) and cross-validation (RMSECV) of each experimental model considering level of detection of each bacterium and the number of components included. Misclassified samples outcome after PLS-DA analysis.

|  |  |  |  |  |  |
| --- | --- | --- | --- | --- | --- |
| Experiment | Detection level (CFU/ml) | Number of components | RMSEC | RMSECV | Misclassified samples |
| *B. subtilis* (V) | 3 x 10<sup>8</sup> | 3 | 0.28 | 0.31 | None |
| *S. enterica* | 50 | 3 | 0.058 | 0.066 | None |
|  | ≤10 | 3 | 0.062 | 0.073 | None |
| *S. haemolyticus* | 50 | 3 | 0.18 | 0.2 | None |
|  | ≤10 | 3 | 0.29 | 0.33 | None |
| *B. subtilis* (S) | 50 | 4 | 0.29 | 0.45 | None |
|  | ≤10 | 4 | 0.28 | 0.63 | None |

## Results and discussion

### Proof-of-concept

We aimed to generate a novel approach using dispersive RS associated with partial least squares discriminant analysis (PLS-DA) to detect bacteria within the DP primary packaging. The first goal was to analyze if the Raman spectra from the bacteria could be distinguished from those related to molecules present in the DP. A *proof-of-concept* was designed to up-concentrate bacteria in a smaller area of the DP container and thus increasing the chance for the laser to find the contaminants. For this purpose, the samples were centrifuged in special plates holding the vials in an inclined upside-down position. This method was efficient in localizing the bacteria along the upper part of one side of the product vial, i.e., close to the neck. To obtain a clearer bacterial spectrum for model training vegetative form of *B. subtilis* were injected into five DP samples with a final concentration of 3 × 10^8^ CFU/ml. Raman spectra from spiked samples with vegetative *B. subtilis* (SS-vBS) and non-injected negative controls (NC) were taken covering the whole area within the vials where the bacterial pellet was localized after centrifugation. Raman spectra after baseline correction and normalization of each group are presented in **Fig. 1A**. Both groups presented similar spectra but with small variations in intensity in peaks related to bacterial-associated organic molecules observed in the wavenumber ranges 700-1800 and 2800-3200 cm^-1^ **(Fig. 1A)**. PLS-DA model was selected due to its power to maximize the inter-class variation and its prediction capacity among samples with unknown degree of within-group variability ^17, 24, 25^. Considering up-till ten components, analysis of variance explanation **(Fig. 1B)**, indicated that three components represented 86.4% of the variation on the dataset. A smaller variation was enclosed in components four and five. Afterwards, any extra component added did not solely explain a significant part of the remaining variation **(Fig. 1B)**. PLS-DA classification plot **(Fig. 1c)** showed that a three-component model was sufficient to discriminate the Raman spectra from SS-vBs and NC. Variation observed among samples, mainly on SS-vBS, could be related to measurements from different locations from these vials. Closer inspection of the loading vectors (LV) showed distinct bacteria-associated peaks, which weighted positively for SS-vBS on the score plot **(Fig. 1d-e)**. SS-vBs received positive scores for peaks on the wavenumbers 786, 814-850 and 1090 cm^-1^ (nucleic acid region), 1000 cm^-1^ (aromatic amino acids), 1060 -1200 cm^-1^ (fatty acids C—C stretching), 1440 cm^-1^ (C—H vibration), and 1656 cm^-1^ (amide I vibration) among the most relevant **(Fig. 1d-e)** ^19, 22, 26, 27^. Some bands located in regions 703-760 and 820-840 cm^-1^ indicating aromatic amino acids can also be related to preservatives (m-cresol and phenol), present in the product formula ^28, 29^. The high wavenumber region 2800-3200 cm^-1^ (C—H stretching region) was kept in the analysis since it has previously been related to major bacterial macromolecules ^26, 30^ that are not present in the DP. LV1 and LV2 presented peaks with negative intensity amplitude in the Raman shift 746, 1000, 1029, 1090, 1 cm^-1^ **(Fig. 1e)**. The latter were important in the NC samples, which received negative scores specially in LV1 **(Fig. 1d-e)** possibly related to the absence of bacterial structures. The negative bump observed in the region at 1300-1450 cm^-1^ in LV1 **(Fig. 1e)** is mainly associated to BO_4_ and SiO_4_ bonds from the primary packaging made of borosilicate glass ^31^.

**Figure 1:**
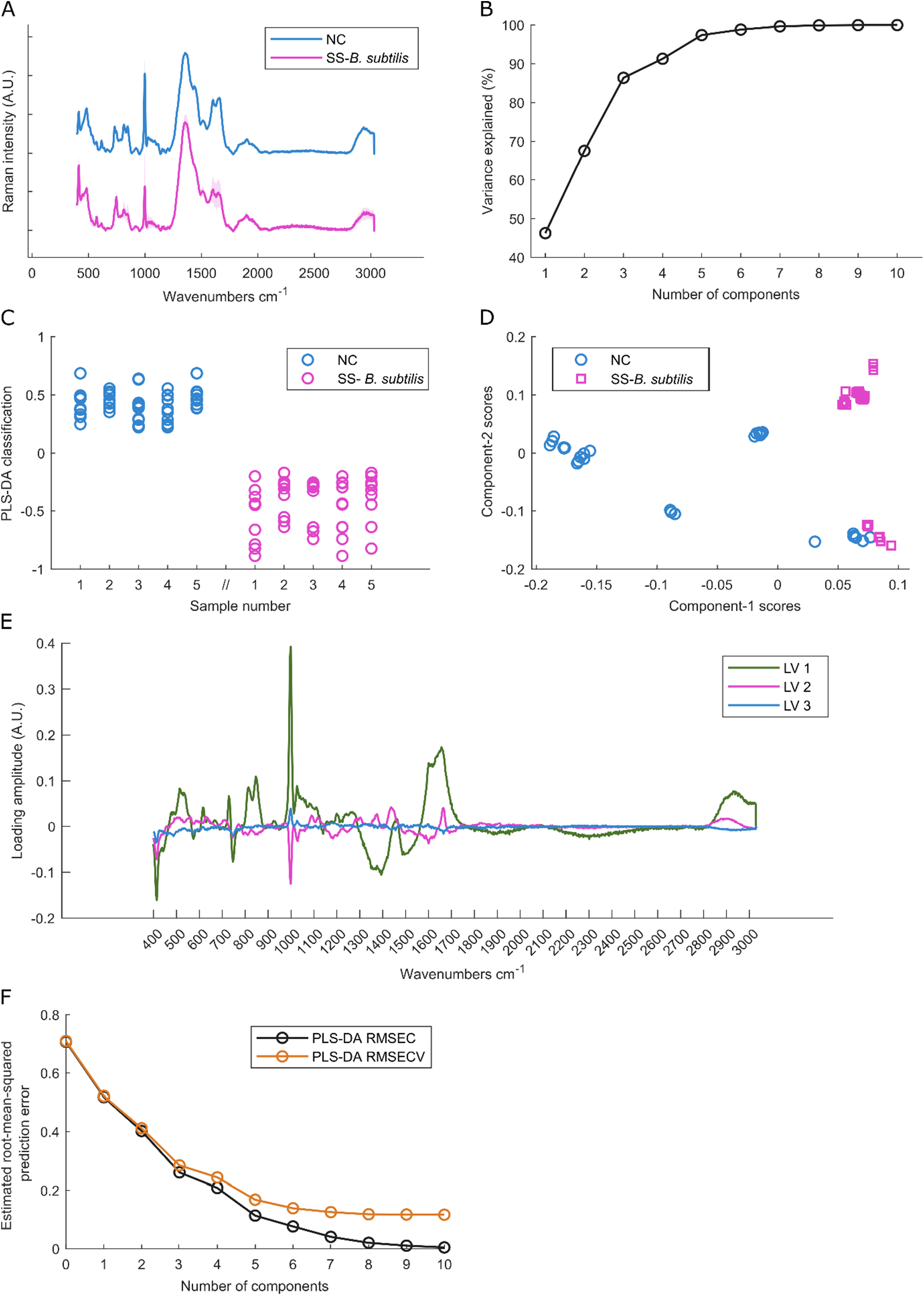
Detection of bacteria in drug product primary package. Raman spectroscopy in combination with partial least squared discriminant analysis (PLS-DA) to distinguish 5 spiked samples with 3 × 10^8^ CFU/ml of vegetative *B. subtilis* (SS-vBs) from 5 negative control (NC) samples. **(A)** Baselined corrected and normalized Raman spectra from SS-vBs (magenta) and NC (blue) represented by data mean (solid line) and ±SD (shadow). **(B)** Variance explained plot of the PLS-DA model in function of the number of components into consideration. **(C)** Classification plot after analysis with a three-component PLS-DA model. Circles represent each measurement of the spiked (magenta) and NC (blue) samples along the vial. **(D)** Score plot of the samples considering scores obtained in component 1 (X-axis) vs components 2 (Y-axis). **(E)** Loading vectors (LV) extracted from the three-component PLS-DA analysis, colors represent each LV1 (green), LV2 (magenta) and LV3 (blue). **(F)** Plot of the estimated root-mean-squared error of calibration (RMSEC, black line) and cross-validation (RMSECV, orange line) in function to the number of components added into the model.

A 10-fold cross-validation was performed to analyze the number of components needed in order to obtain the best performance of classification through the root mean squared error of cross-validation (RMSECV) and comparing with the overall fitting by RMSE of calibration (RMESC) **(Fig. 1f)**. Based on RMSECV **(Fig. 1f – orange line)**, five components were the limit for error reduction and onwards we had a diminishing in return. However, three components were enough to perform without misclassification reducing the risk of overfitting and making the model more robust in terms of classifying future samples that are different to the training dataset. In the **Table 1** are summarized all RMSEC and RMSECV values obtained for each experiment as well as their respective number of components considered, and the outcome of samples misclassified.

In our system we used 785 nm visible light, since water has a relatively low absorption at this wavelength while obtaining a relatively strong signal from bacteria in comparison to other wavelengths ^14^. In addition, the 785 nm laser presented less fluorescence interference and improved signal-to-noise ratio compared to other visible light spectra, such as a 532 nm laser ^32-34^. Supporting this choice, previous studies have shown great robustness of 785 nm in detecting wide spectrum of both Gram-positive and Gram-negative bacteria ^35^. In summary, this concept adding a special centrifugation of vials could improve the bacterial detection with RS in DP. Moreover, the PLS-DA approach allowed us to maximize the inter-class differences and therefore discriminate the Raman signal of bacteria suspended in DP from the DP alone within the primary packaging.

### Detecting low number of vegetative bacteria within primary package

Major challenges encountered when reaching low CFU detection limits with RS are: (i) to discriminate bacterial signal from other organic compounds present in the DP without compromising the model accuracy, (ii) to detect bacteria in concentrations lower than 100 CFU/ml keeping the cells viability for future identification procedures, and (iii) to detect bacteria and discard the primary packaging signal while keeping the integrity of the DP primary container ^14, 36-38^. Therefore, we challenged our RS-PLS-DA model to detect vials containing below 100 CFU/ml of two important yet different bacteria: *Salmonella enterica* (rod-shaped gram-negative) and *Staphylococcus haemolyticus* (coconut-shaped gram-positive). It is important to mention that we did not focus on distinguishing between the bacterial species but on confirming the presence or absence of contaminants with high reproducibility.

Following the same procedure of spiking and centrifugating as explained before, we prepared samples with each species reaching a final concentration of 50 CFU/ml and ≤10 CFU/ml. Raman spectra were obtained from SS containing 50 CFU *S. enterica* (50-SE), ≤10 CFU *S. enterica* (10-SE), 50 CFU *S. haemolyticus* (50-SH) and ≤10 CFU *S. haemolyticus* (10-SH) and compared against NC **(Fig. 2)**. In SS, given the inclined centrifugation of the vials we expected to obtain a mixture of positive bacterial signal from the first half of the vial and negative onwards. After Raman analysis the samples were cultured to compare results with the ones obtained from RS. Explorative analysis of the training dataset was performed using the same PLS-DA model and corroborate if consistent output variables were obtained i.e., discrimination between samples with and without bacteria. PLS-DA classification and score plots of loading vectors are shown in **Fig. 2a-h**. A three-component model showed a well separation of samples with bacteria and NC of both concentrations of *S. enterica* and *S. haemolyticus* **(Fig. 2a,c,e,g)**. Both experiments with *S. enterica*, 50-SE and 10-SE, received a better discrimination in terms of inter-class distance (Y-axis) from NC group in comparison to *S. haemolyticus* samples **(Fig. 2a, c)**. However, score plots of LV1 and LV2 showed a correct discrimination of SS and NC training dataset clusters even at two-component model for all bacterial specie and concentration analyzed **(Fig. 2 b,d,f,h)**. On the score plots, groups representing SS with 50-SE and 10-SE appeared as two separated subclusters with different weight scores for LV1 and LV2 **(Fig. 2b,d)**. This intra-group gap reflected the measurements from areas with bacterial signal and without bacteria from centrifuged samples bearing bacteria. Concerning *S. haemolyticus*, in both 50-SH and 10-SH groups the inter-class separation was less prominent compared to *Salmonella* and observations from the negative areas appeared closer to the sterile group **(Fig. 2e,g)**. This was also demonstrated in the score plot where three sub-clusters were defined for SS with 50-SH and 10-SH, indicating more variability among these observations **(Fig. 2f,h)**. In these plots it is also observed that some observations from SS at both concentrations scored similar to sterile groups. Despite that, no misclassification of training dataset was obtained **(Table 1)**. The difference observed in term of greater discrimination for SS with *S. enterica* in comparison to *S. haemolyticus* could be related to bacterial composition, size and shape being reflected in Raman spectra peaks intensity ^35^. Supporting this idea, experiments detecting both 50-SE and 10-SE the RMSECV obtained was remarkable lower under three-components compared to detection of 50-SH and 10-SH (**Table 1)**. Bacteria culture was representative of Raman results obtaining growth of bacteria colonies for all SS evaluated and no growth on the NC (data not shown)

**Figure 2:**
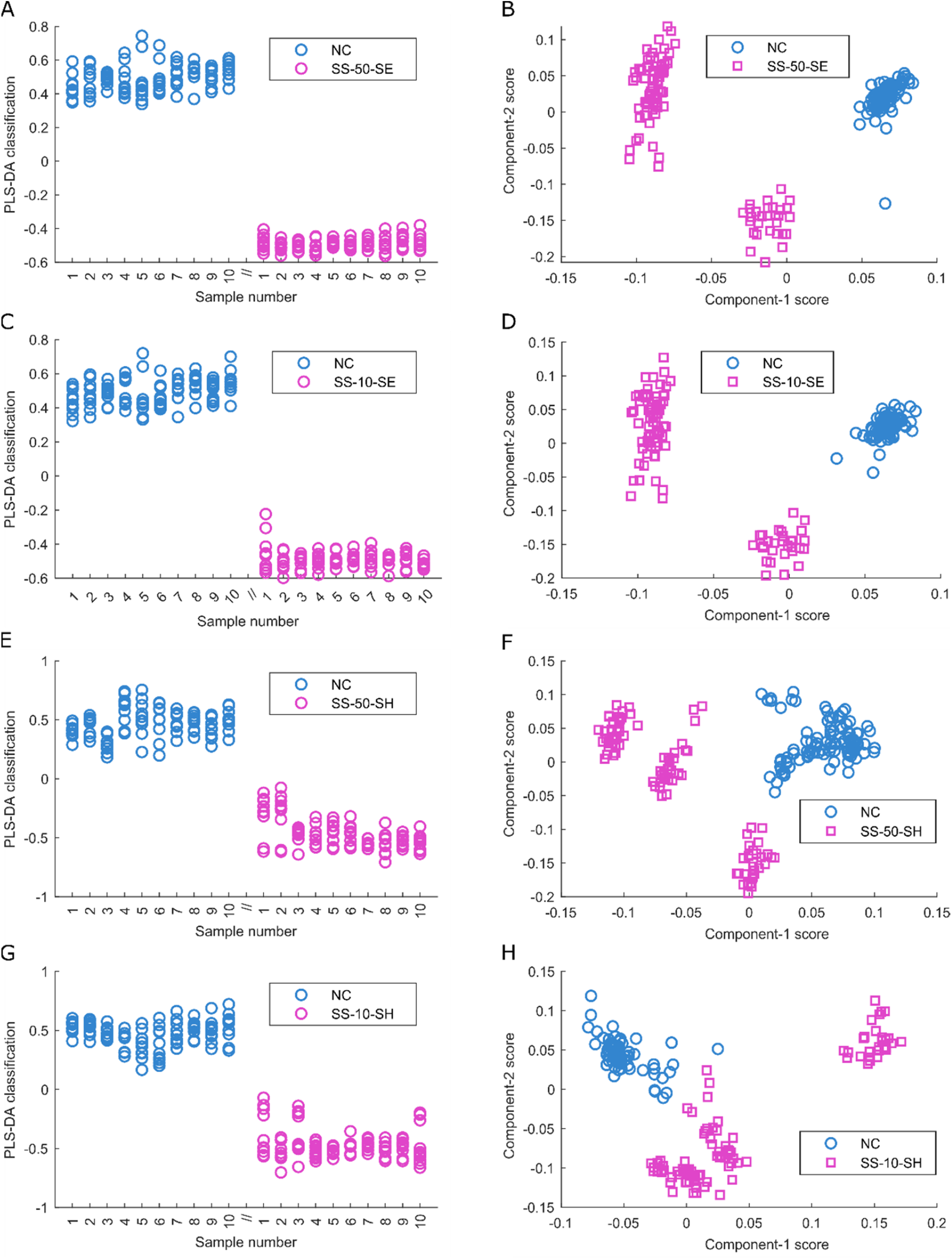
Detection of low bacterial concentrations in vegetative forms. Raman spectra from 10 spiked samples (SS) and 10 negative controls (NC) were collected and analyzed using a three-component partial least squared with discriminant analysis (PLS-DA) model. Classification and score plots obtained in components number 1 and 2 of **(A-B)** SS with 50 CFU/ml of *S. enterica* (SS-50-SE); **(C-D)** SS with 10 CFU/ml of *S. enterica* (SS-10-SE); **(E-F)** SS with 50 CFU/ml of *S. haemolyticus* (SS-50-SH); **(G-H)** SS with 10 CFU/ml of *S. haemolyticus* (SS-10-SH).

### Low LOD of bacterial spore detection

Some bacteria relevant for pharmaceutical production have the ability to produce spores as a survival mechanism in a hostile environment. A method used for quality control of pharmaceutical presentations required to be sterile should also be able to detect spores, especially given the increased likelihood of bacteria in spore form surviving unfavorable conditions for extended time periods and reaching the patient. Hence, we challenged our RS-PLS-DA model to detect low concentrations of *B. subtilis* spores (BSs). We spiked vials with 50 CFU and ≤10 CFU/vial of *B. subtilis* spores (50-BSs and 10-BSs respectively) and then analyzed with RS and compared to NC. In **Figures 3a** and **3c** is presented the PLS-DA classification for both 50-BSs and 10-BSs respectively. The model correctly discriminated the samples even though in both concentrations the variation between classes was less prominent in comparison to the case with vegetative cells **(Fig. 2)**. The less prominent discrimination was supported by the intra-sample variation as well as the limited inter-class separation observed on the Y-axis **(Fig. 3a, c)**.

**Figure 3:**
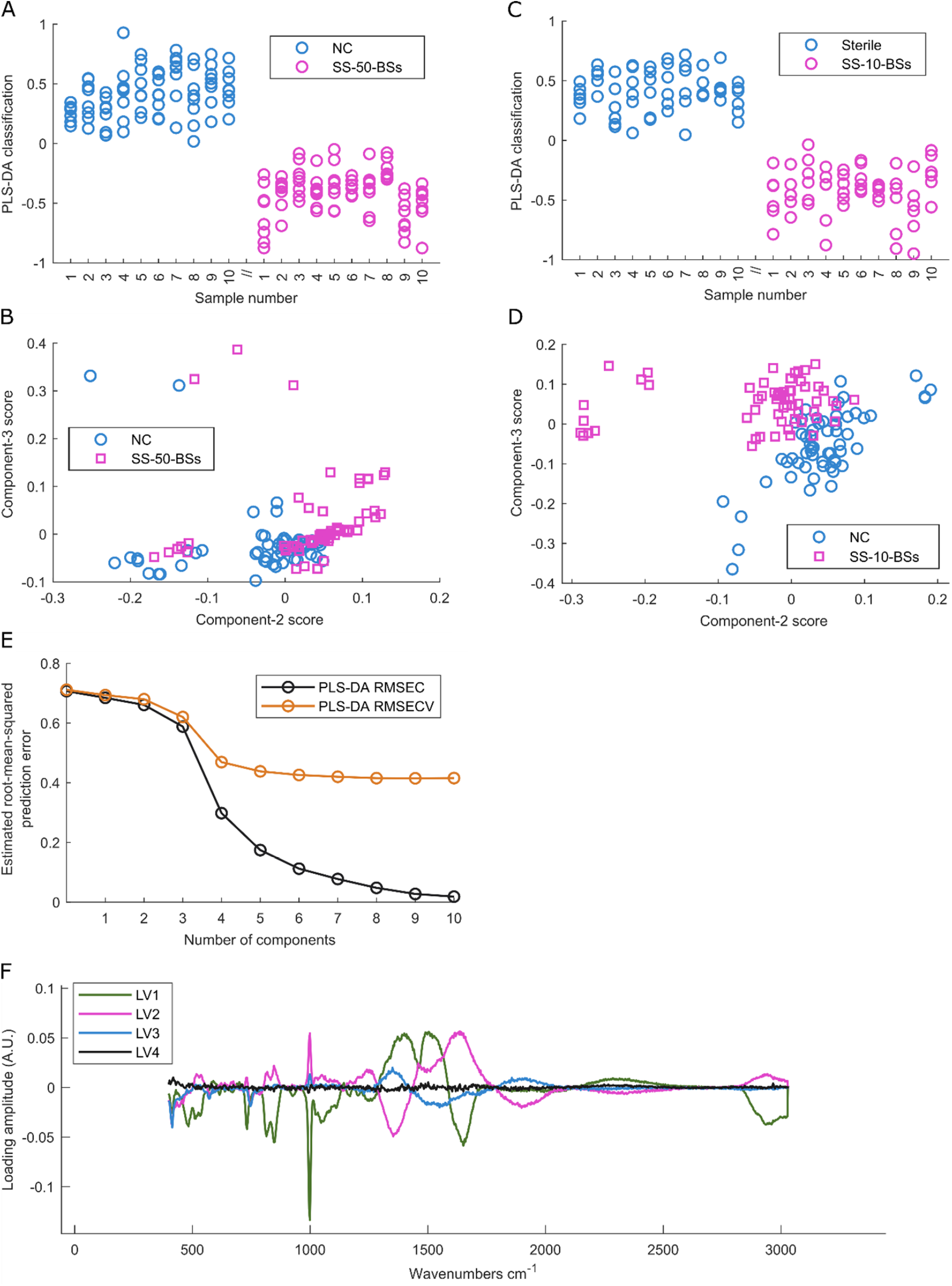
Detection of spores. Raman spectra from 10 spiked samples (SS, magenta) and 10 negative controls (NC, blue) were collected and analyzed using a four-component partial least squared with discriminant analysis (PLS-DA) model. Classification and score plots obtained in components number 2 and 3 of **(A-B)** SS with 50 CFU/ml of *B. subtilis* spores (SS-50-BSs); **(C-D)** SS with 10 CFU/ml of *B. subtilis* spores (SS-10-BSs). **(E)** Plot of the estimated root-mean-squared error of calibration (RMSEC, black line) and cross-validation (RMSECV, orange line) in function to the number of components added into the model. **(F)** Loading vectors (LV) extracted from the four-component PLS-DA analysis to visualize peaks related to spore discrimination, colors represent each LV1 (green), LV2 (magenta), LV3 (blue) and LV4 (black).

For spore detection at very low cell numbers, four components were needed to reach a complete group separation instead of the two-component needed for vegetative cells. This was observed on the score plots were overlapping among the observation was remanent when comparing LV2 vs LV3 **(Fig 3b, d)**. In addition, in the RMSECV plot an evident improving step in error reduction could be seen when jumping from three components to four **(Fig. 3e)**. In a general perspective it is clearly observed that RMSEC and RMSECV obtained in both spore concentrations were higher compared to the errors observed in vegetative bacterial forms (**Table 1)**. Loading plot of LV1-4 showed weak but clear Raman signal related to spores which was mainly supported by the peaks at 786, 814-850 and 1090 cm^-1^ (nucleic acid), 1464 (lipids), 1579, 1665 cm^-1^ (proteins) and 2800-3200 cm^-1^ (C—H) ^30, 39, 40^ **(Fig. 3f)**. At this spore concentration, previously reported specific peaks related to *Bacillus sp*. spores in the wavenumbers 824, 1017 1395 and 1579 cm^-1 30, 39, 41, 42^ were difficult to spot given their weak signal in comparison to other peaks. **(Fig. 3f)**. The latter is possibly related to different factors, first that the model was built and trained based on vegetative forms which are phenotypically different to spore form ^19, 26^. The second reason could be associated to the low working concentration making it more difficult to visualize spore-specific peaks in comparison to larger and more predominant macromolecules, which already had a weak signal in the LVs. Spores are around one-third of the size of a vegetative cell ^43, 44^, so it is more difficult to obtain a clear Raman signal from them and thus also to detect low numbers of spores. Is important to highlight that even if most authors consider wavenumbers from 400 to 1800 cm^-1^ as the most active region for bacterial Raman signal, with the spores we should also consider the high wavenumber region 2800-3000 cm^-1^. The latter region is related to predominant spore components such as fatty acids, carbohydrates, and proteins, which are packed together with the genetic material ^19, 26^. These results together demonstrate that our RS-PLS-DA model is not affected by bacterial phenotypic features but instead focuses on common components of them.

### Model validation

Cross-validation of independent samples is crucial to demonstrate the ability of the model to discriminate new samples with unknown contamination. For this purpose, we performed a 10-fold leave-one-out cross-validation analysis using our RS-PLS-DA model and a new independent set of samples. Contamination with *B. subtilis* spores were used given they were the most challenging case concerning detection of low numbers. Two datasets were used for cross-validation: (i) a training data set using samples from **Fig. 3c** and (ii) an independent (validation) set with 20 new samples comprising ten vials with ≤10 CFU-BSs and ten sterile vials. After Raman analysis, samples were cultured to compare results with the ones obtained by RS-PLS-DA. Classification of training (stars) and validation (triangles) datasets from cross-validated data are presented in **Fig. 4a**. Both sterile (blue triangles) and contaminated (red triangles) validation groups were discriminated with 99% specificity and 98.3% sensitivity, establishing a hard decision cut-off in zero (dotted line) **(Fig 4a)**. Validation samples dataset presented larger intra-group variations compared to the training group. This represented a realistic scenario of new samples being compared with the training dataset building the PLS-DA model. RMSE of validation groups (RMSEV) **(Fig. 4b, blue line)** showed that an improvement could be obtained up till five components considered. This reduction was not seen by analyzing RMSECV of the independent validation (black line) where after the third component any additional component did not reduce the prediction error **(Fig 4b)**. The latter is important when we take into count that the results showed in **Fig. 3**, indicated that four components were needed in order to discriminate 50 and 10 CFU of BSs from NC without misclassification **(Table 1)**. Therefore, using less components some samples crossed the decision line generating some false positive and false negative observation which impacted on the accuracy obtained **(Fig 4a)**.

**Figure 4:**
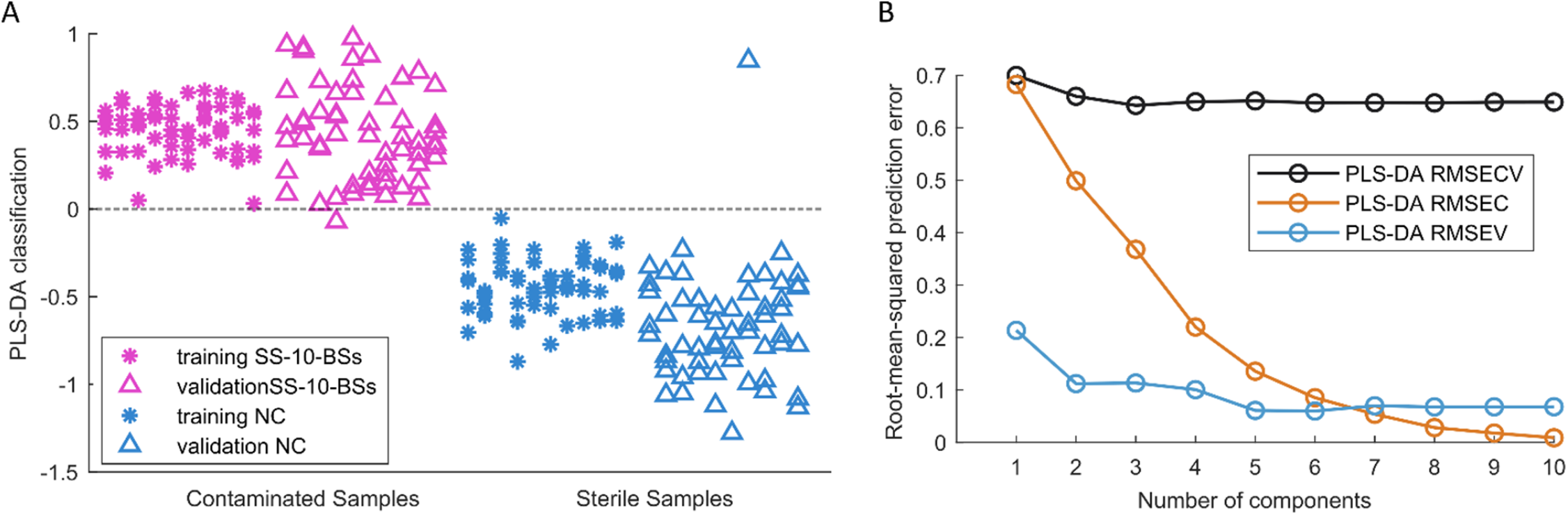
Model validation. Independent validation using partial least squared discriminant analysis (PLS-DA) and 10-fold cross-validation. **(A)** Classification of 10 spiked samples with 10 CFU/ml of *B. subtilis* spores (SS-10-BSs) and 10 negative controls (NC) were used as training dataset (magenta and blue stars respectively). 10 new SS-10-BSs and 10 NC samples were used as independent test set (magenta and blue triangles respectively). **(B)** Plot of the root-mean-squared error of calibration (RMSEC, orange), cross-validation (RMSECV, black line) and validation (RMSEV, blue line) in function to the number of components added into the model.

In this work, we obtained similar values in terms of accuracy in comparison to previous reports showing sensitivities and specificities around 95-99% using RS coupled with multivariate analysis models. However, some approaches do not specify the limit of detection reached ^45^ or they presented theoretical detection limits below 100 CFU/ml based on RMSECV values obtained on the calibration and not in physical measurements ^9^. Another common drawback is to ensure reproducibility of the results using a single model to detect samples with less than 10 CFU/ml or when cross-comparing among different species ^22, 26, 35, 37^. In addition, our results showed a great performance about the grade of robustness including three highly different bacteria species and bacterial spores. Another important feature is that we used a non-invasive method to reach low LOD which allowed us to test the DP within its container. The latter is an improvement of previous concepts involving complex RS systems to obtain low LOD involving procedures breaching the integrity of the DP samples such as needing external amplifier particles and/or dried samples on metallic surfaces ^37, 46^. In this sense, sample handling that requires breaching of the barrier presented by the original container also poses a risk of obtaining secondary sample contaminations thereby giving a false positive result. Time consumption is also a drawback when analyzing samples with RS concepts involving several steps, especially if the final application is in pharmaceutical industry. Even though LOD below 10^3^ CFU/ml could be secured, these kinds of multiple-step and advance systems need many hours to analyze a single small sample ^37^ On the contrary, we only needed around 3 minutes to evaluate each sample containing 3 ml of product with RS after centrifugation, and the evaluation is done without breaching the barrier of the original sample container. Further analyses are needed to deepen in understanding the limits of RS technique for application in contamination control such as detection of cell debris, death cells and other relevant scenarios that may cause ambiguous results.

## Conclusions

In the present study we evaluated a fast and non-invasive approach to discriminate pharmaceutical products vials containing low numbers of bacteria from sterile ones using dispersive Raman spectroscopy in association with PLS-DA. The RS-PLS-DA concept was challenged to detect bacterial species suspended in drug product within its primary package. Three highly different bacteria including *B. subtilis* spores were detected without breaching the DP vial. This rapid and simple concept innovates in its effective way to localize the contaminants in a smaller area within the product vial in order to secure laser detection. Our results showed a successful discrimination when detecting vegetative cells and spores at the very low concentration of 10 CFU/ml, even in the presence of other organic molecules from the product formula in the intact DP primary container. True independent validation showed an outstanding performance with high sensitivity and specificity. In summary, we provide a feasible approach using RS in association with PLS-DA to detect extremely low numbers of cells or spores with high accuracy and reproducibility without compromising the robustness of the method. These results support Raman spectroscopy as a promising biotechnological tool suitable for bioburden test in quality control of pharmaceutical industry.

## Author contributions

G.R.A. conceived and performed the experiments, analyzed the data, discussed the results, and wrote the manuscript. W.A.R. and H.M.A.B. performed the data processing script and analyzed the data. S.B. provided project design ideas and discussed the results. B.E and S.A provided expertise for experimental design, discussed the results, and provided feedback. O.K.E and Q.H. provided expertise and feedback. A.E.C. coordinated the project and provided expertise, feedback. All authors read and approved the final version of the manuscript.

## Acknowledgment

Villum young investigator given to A.E.C (19105). Novo Nordisk A/S to A.E.C., for financial support for reagents acquisition.

